# Identification of ESCRT-III like protein sequences in yeast

**DOI:** 10.1101/2024.03.07.583924

**Authors:** Thomas Brune, Ralf Kölling

## Abstract

Endosomal sorting complex required for transport (ESCRT-III) is a membrane remodeling complex involved in a large number of cellular processes. It appears to perform an essential function in eukaryotes, since to date no eukaryotic organism completely devoid of ESCRT-III has been found. Yet, yeast cells with a deletion of all eight known ESCRT-III genes are viable. We therefore searched for new, previously undiscovered ESCRT-III like proteins in yeast. HHPred uncovered several proteins with similarity to Snf7. The similarity was mostly restricted to the *α*1-*α*2 hairpin region of Snf7. A conserved pattern of amino acids was detected in this region. The protein encoded by ORF YPL199c strikingly resembled Snf7 in its secondary structure. Since this protein could be the ninth member of the ESCRT-III family in yeast, we called it Nbr9 (“number nine”). Nbr9 is palmitoylated and localizes to the plasma membrane. In contrast to other palmitoylated proteins, it is not associated with lipid rafts. When *NBR9* was deleted in the octuple ESCRT-III deletion background, the yeast cells were still viable. However, despite a number of experiments, we do not have evidence at present that Nbr9 is part of an alternative ESCRT-III complex.

## Introduction

Endosomal sorting complex required for transport (ESCRT-III) is a membrane remodeling factor involved in many different cellular processes like intraluminal vesicle formation at endosomes, cytokinesis, HIV-budding, plasma membrane and nuclear membrane repair, nuclear envelope resealing after mitosis, autophagy or neuron pruning (Hurley 2015; McCullough, Frost et al. 2018; Gatta and Carlton 2019; Vietri, Radulovic et al. 2020). There are twelve members of the ESCRT-III protein family in human cells and eight in yeast. In yeast, where these proteins were first identified, only four of the proteins (Snf7, Vps2, Vps20 and Vps24) are considered to be part of ESCRT-III, while the remaining members of the protein family (Chm7, Did2, Ist1 and Mos10/Vps60) are called ESCRT-III associated proteins (Babst, Katzmann et al. 2002; Azmi, Davies et al. 2008; Dimaano, Jones et al. 2008; Rue, Mattei et al. 2008). However, this distinction does not appear to be justified, since all ESCRT-III proteins can team up in various combinations during different cellular processes. For instance, evidence has been presented in yeast for the existence of at least two alternative ESCRT-III complexes, which are based on Chm7 or Mos10, respectively (Bauer, Brune et al. 2015; Webster, Thaller et al. 2016; Gu, LaJoie et al. 2017; Alsleben and Kölling 2022; Pfitzner, Zivkovic et al. 2023).

All ESCRT-III proteins have the same basic structure, as exemplified by human CHMP3 (Muziol, Pineda-Molina et al. 2006). The ESCRT-III unit consists of six *α*-helices. The first two helices fold into a 70 Å long hairpin that together with two short helices forms a four-helical bundle. The last two helices are part of the largely disordered C-terminal region and are not represented in the crystal structures. The ESCRT-III proteins can exist in an open and in a closed conformation (Bajorek, Schubert et al. 2009). In the closed conformation the *α*5-helix folds back onto the *α*1-*α*2 hairpin, which prevents filament formation. Filament formation seems to be a general property of all ESCRT-III proteins. Filament formation is promoted by the open conformation, where the interaction between the *α*1-*α*2 hairpin and the *α*5-helix is released and *α*2now forms a long, extended helix with *α*3 (Tang, Henne et al. 2015). Although ESCRT-III proteins usually exist in their open conformation in filaments, a filament structure has been described, where IST1 in its closed conformation forms a co-filament with CHMP1B in its open conformation (McCullough, Clippinger et al. 2015).

ESCRT-III is a hallmark of eukaryotic life. Although not all eukaryotic organisms contain the whole complement of ESCRT-III proteins, to date no eukaryotic organism has been identified that is completely devoid of ESCRT-III (Leung, Dacks et al. 2008). In mammals ESCRT-III seems to be essential for life. For instance, it has been shown that downregulation of CHMP5 is embryonically lethal in mice (Shim, Xiao et al. 2006).We deleted all eight ESCRT-III genes in yeast and obtained a yeast strain that is viable and grows like wildtype at ambient temperature (Brune, Kunze-Schumacher et al. 2019). One reason for this unexpected result could be that there are additional members of the ESCRT-III family in yeast that have not been identified yet. Here we present the identification of new putative ESCRT-III like sequences in yeast. One these proteins encoded by yeast ORF YPL199C strikingly resembles the ESCRT-III protein Snf7 in its secondary structure. This protein, which we call Nbr9 (“number nine”), could be a new member of the ESCRT-III family in yeast.

## Materials and methods

### Media, yeast strains and plasmids

Yeast cells were grown in YPD medium (1 % yeast extract, 2 % peptone, 2 % glucose) or in SD/CAS medium (0.67 % yeast nitrogen base, 1 % casamino acids, 2 % glucose, 50 mg/l uracil and tryptophan). The yeast strains RKY3021 (NBR9-3HA::HIS3) and RKY3417 (NBR9-ymNeongreen::HIS3) were derived from JD52 (J. Dohmen, Cologne, Germany) by the integration of PCR-cassettes into the yeast genome (Longtine, McKenzie et al. 1998). The CEN/ARS plasmid with *TRP1* marker pRK1883 carries an *NBR9-sfGFP* fusion under the control of the *HXT7* promoter. The cysteine codons 233 and 235 of *NBR9* were mutagenized to serine codons by QuikChange mutagenesis (Agilent, Waldbronn, Germany) to give plasmid pRK1891.

### Fluorescence microscopy

Yeast cells were grown overnight to exponential phase in SD/CAS medium. The yeast cell suspension was applied to concanavalin A coated slides and imaged with a Zeiss Axio-Imager M1 fluorescence microscope equipped with an AxioCam MRm camera (Zeiss, Göttingen, Germany). Images were acquired with the Axiovision Software and processed with Photoshop Elements.

### Flotation

Yeast cells were grown overnight to exponential phase in YPD medium. 10 OD_600_ of cells were harvested, washed in 10 mM NaN_3_, resuspended in 100 μl TNE-buffer (50 mM Tris-Cl pH 7.4, 150 mM NaCl, 5 mM EDTA) with protease inhibitors and lysed by glass beading for 5 min at 4°C. After the addition of 150 μl of TNE, the cell extract was spun at 500 g for 5 min to remove cell debris. 125 μl of the supernatant were mixed with 250 μl 60 % OptiPrep (Merck, Darmstadt, Germany). This mixture was overlaid with 30 % OptiPrep (diluted 1:2 with TNE) and with 100 μl TNE. The step gradient was centrifuged for 2 h at 100,000 g. Six 180 μl fractions were collected from the top of the gradient, mixed with one volume of 2-times SDS sample buffer and heated to 95° for 5 min.

## Results and Discussion

Previously, we generated a yeast strain with a deletion of all eight known ESCRT-III genes (Brune, Kunze-Schumacher et al. 2019). ESCRT-III activity is essential in mammalian cells (Shim, Xiao et al. 2006). Thus, it came as a surprise that our octuple deletion strain was viable. This prompted us to search for new, previously undiscovered members of the ESCRT-III family in yeast. For this we used HHPred, which is more sensitive than BLAST or PSI-BLAST and is able to detect remote homologs (Zimmermann, Stephens et al. 2018). With the ESCRT-III protein Snf7 as a query the output shown in Fig. S1 was obtained. The seven top hits were yeast ESCRT-III proteins, which were predicted with a probability close to 100 %. The atypical ESCRT-III protein Ist1 was number fifteen on the list (probability: 63.56 %, E-value: 11).

The five proteins detected by HHpred between the ESCRT-III proteins on top of the list and Ist1 (some are listed multiple times) were analyzed in more detail for their similarity to ESCRT-III proteins. The secondary structure was examined by Jpred (Drozdetskiy, Cole et al. 2015) and a structure prediction was made with AlphaFold (Varadi, Anyango et al. 2022) (Fig. 1). The regions of similarity with Snf7, the corresponding regions in Snf7 and the HHPred confidence scores are summarized in Tab. 1. All detected proteins contain extended *α*-helical regions. For the FG-nucleoporin Nsp1 the similarity to Snf7 extends to the whole *α*1-*α*4 core.

**Table 1.** HHPred regions of similarity between target proteins and Snf7.

| Protein | Size<br>aa | Region of<br>similarity<sup>1</sup> | Similarity to<br>Snf7<sup>1</sup> | Snf7<br>region | Probability<br>(%) | E-value |
| --- | --- | --- | --- | --- | --- | --- |
| Nsp1 | 823 | 635-806 | 17-180 | $\alpha 1$-$\alpha 4$ | 93.90 | 0.8 |
| Nnf2 | 936 | 730-889 | 19-178 | $\alpha 1$-$\alpha 4$ | 87.14 | 3.9 |
| Jac1 | 184 | 121-174 | 30-83 | $\alpha 1$-$\alpha 2$ | 73.45 | 4.8 |
| Gta1 | 956 | 288-363 | 19-96 | $\alpha 1$-$\alpha 2$ | 65.84 | 18 |
| Nbr9 | 240 | 27-82 | 31-86 | $\alpha 1$-$\alpha 2$ | 64.85 | 9.3 |

<sup>1</sup>amino acid position

**Figure 1.**
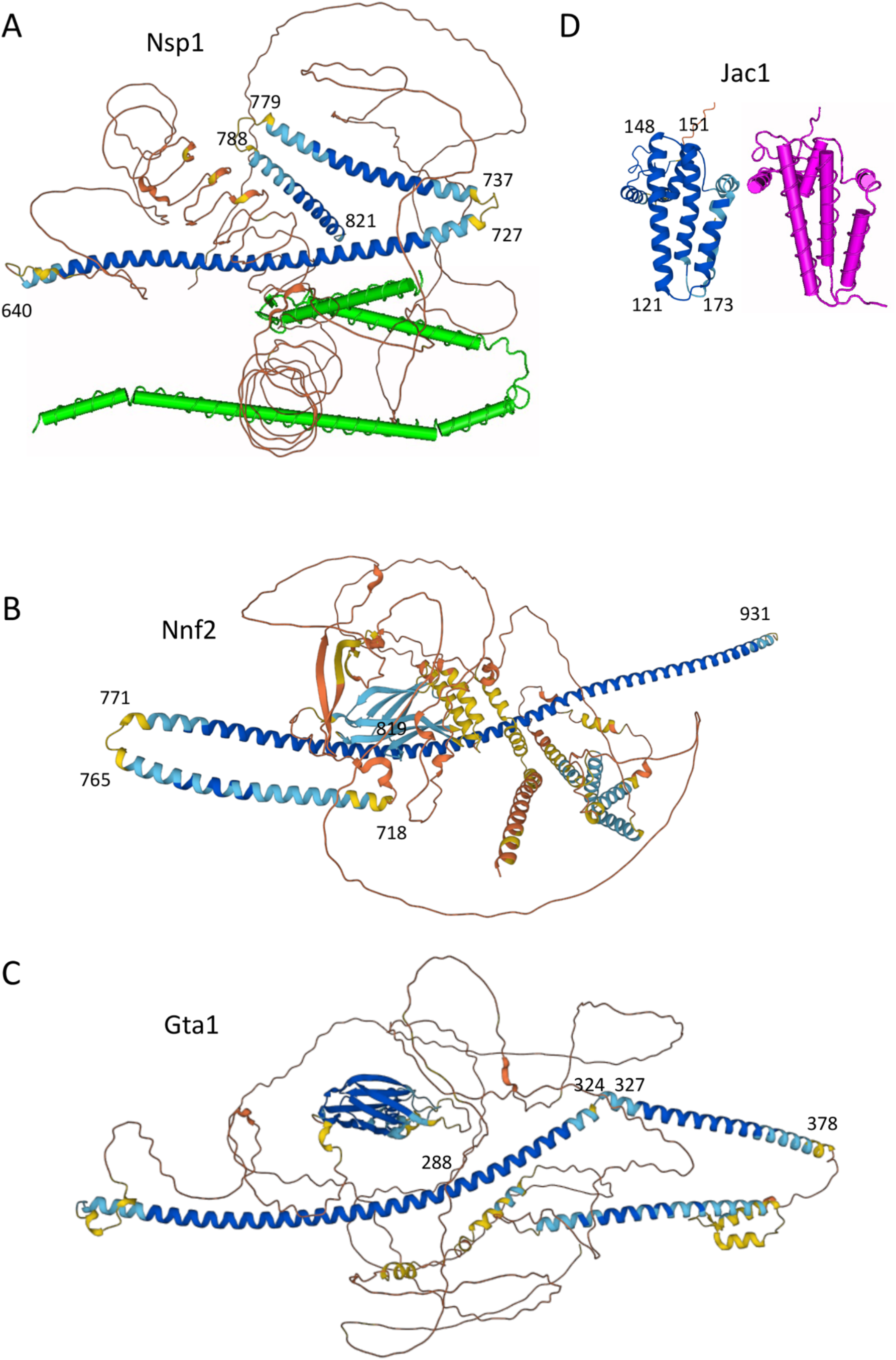
Structure prediction by AlphaFold. (A) Nsp1, top: AlphaFold prediction, bottom: cryo-EM structure of the *α*-helical region of Nsp1 (green) (Li, Chen et al. 2022), (B) Nnf2, (C) Gta1/Yel043w, (D) Jac1, left: AlphaFold prediction, right: 3D-structure (magenta) (Ciesielski, Schilke et al. 2012). Numbers mark the sequence positions of the regions similar to Snf7 (Tab. 1).

AlphaFold predicts a long rod-like *α*-helix corresponding to *α*1-*α*2 of Snf7 followed by the *α*3 and *α*4 counterparts. In contrast to Snf7 the *α*1-*α*2 helices do not form a hairpin structure, but are extended. Nsp1 is part of the nuclear pore complex, for which a cryo-EM structure is available (Li, Chen et al. 2022). This opens up the opportunity to validate the AlphaFold prediction. The cryo-EM structure of Nsp1 perfectly matched the AlphaFold prediction (Fig. 1A). The connection between Nsp1 and ESCRT-III could be more than accidental. ESCRT-III is involved in the formation of intraluminal vesicles (ILVs) at endosomes. The forming ILV bulges away from the cytosol and forms a structure, which is topologically similar to the membrane tube between outer and inner nuclear membrane at the nuclear pore. The Nnf2 protein, which is supposed to be associated with RNA polymerases (Briand, Navarro et al. 2001), also has a long *α*-helical region similar to the *α*1-*α*4 core of Snf7. But in contrast to Nsp1, the region equivalent to *α*1-*α*2 forms a hairpin, like in Snf7 (Fig. 1B).

The Gta1 protein (ORF YEL043w), proposed to function in Golgi vesicle trafficking (Mattiazzi Usaj, Sahin et al. 2020), has a long *α*-helical region, where only a part in the middle of this extended region bears similarity to *α*1-*α*2 of Snf7 (Fig. 1C). The putative J-type chaperone Jac1 is different from the previous proteins, in that it forms a compact *α*-helical structure. Two of its helices are similar to *α*1-*α*2 of Snf7 (Fig. 1D). For Jac1 a 3D-structure is available (Ciesielski, Schilke et al. 2012) and again the structure was perfectly predicted by AlphaFold. This underlines the predictive power of AlphaFold. The remaining candidate protein, Ypl199c, will be described in more detail below.

The question arises, whether the detection of the Snf7 target proteins by HHPred was purely based on the presence of extended *α*-helical regions. For this reason, we looked for similarities in the detected helices. An alignment of the regions of the five candidate proteins corresponding to the *α*1-*α*2 region of Snf7 was generated with ClustalX. The alignment revealed a pattern of stripes of hydrophobic and positively or negatively charged amino acids (Fig. 2). In the first half of *α*1 there are three stripes of positively charged amino acids, while negatively charged amino acids are concentrated at three positions in the second half of *α*2. It is tempting to speculated that this distribution of amino acids could help to stabilize the *α*1-*α*2 hairpin by electrostatic interactions. Even in the cases where no hair-pin is predicted alternative conformations with a hair-pin could exist *in vivo*. On top of this, a conserved pattern of hydrophobic amino acids was observed. We considered the possibility that this pattern could be related to the formation of coiled-coils, even though the spacing does not conform to a heptad repeat pattern as observed in coiled-coil proteins. In our initial identification of part of the ESCRT-III protein family we noted the propensity of the proteins to form coiled-coils (Kranz, Kinner et al. 2001). The coiled-coil forming ability of the regions similar to Snf7 was analyzed with PCOILS (Zimmermann, Stephens et al. 2018) (Fig. S2). Coiled-coils were predicted for the *α*1, *α*2 and *α*5 helices of Snf7, but for the other proteins no consistent coiled-coil pattern was observed. In the case of Nnf2 and Gta1 all helices similar to Snf7 had the potential to form coiled-coils. But, for Nsp1 and Jac1 no coiled-coils were predicted in the *α*1-*α*2 region. For Ypl199c only the *α*1-helix had a propensity to form coiled-coils. Thus, there is no clear correlation between the pattern of hydrophobic amino acids in the *α*1-*α*2 region and a coiled-coil pattern. Also, no typical coiled-coil proteins were detected by HHPred with Snf7 as a query. When the *α*1-*α*2 region of Snf7 was replaced by *α*-helical regions of the coiled-coil protein Mlp1 (Kölling, Nguyen et al. 1993), a large number of coiled-coil proteins were pulled out by HHPred (not shown). Thus, HHPred clearly detects a distinct pattern of amino acids in the Snf7 targets, different from a coiled-coil pattern.

**Figure 2.**
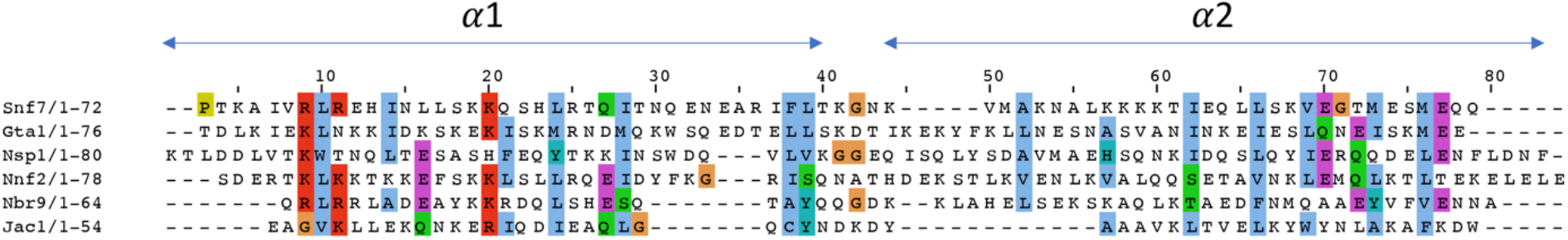
Alignment of the regions similar to the *α*1-*α*2 region of Snf7. The five protein sequences with similarity to the *α*1-*α*2 region of Snf7 were aligned with Clustal X. The amino acid residues are colored according to the Clustal X default coloring scheme (Fig. S2). The *α*1 and *α*2 helix regions of Snf7 are indicated on top of the diagram.

In the following we took a closer look at the remaining Snf7 target Ypl199c. The analysis of the secondary structure revealed a striking similarity to Snf7 (Fig. 3). Thus, Ypl199c could be a new member of the ESCRT-III family. Since it would be the ninth member of the ESCRT-III family in yeast, we named the protein Nbr9 (“Number nine”). Nbr9 and Snf7 both consist of 240 amino acids and three of the six *α*-helices of Snf7 (*α*1, *α*2 and *α*6) are perfectly conserved with respect to length and position in Nbr9 (Tab. 2). The similarity detected by HHPred lies in the *α*1-*α*2 hairpin of Snf7. The main difference between Snf7 and Nbr9 is that Nbr9 contains a “small MutS-related (SMR)” domain (Fukui and Kuramitsu 2011) in the middle of the protein.

**Figure 3.**
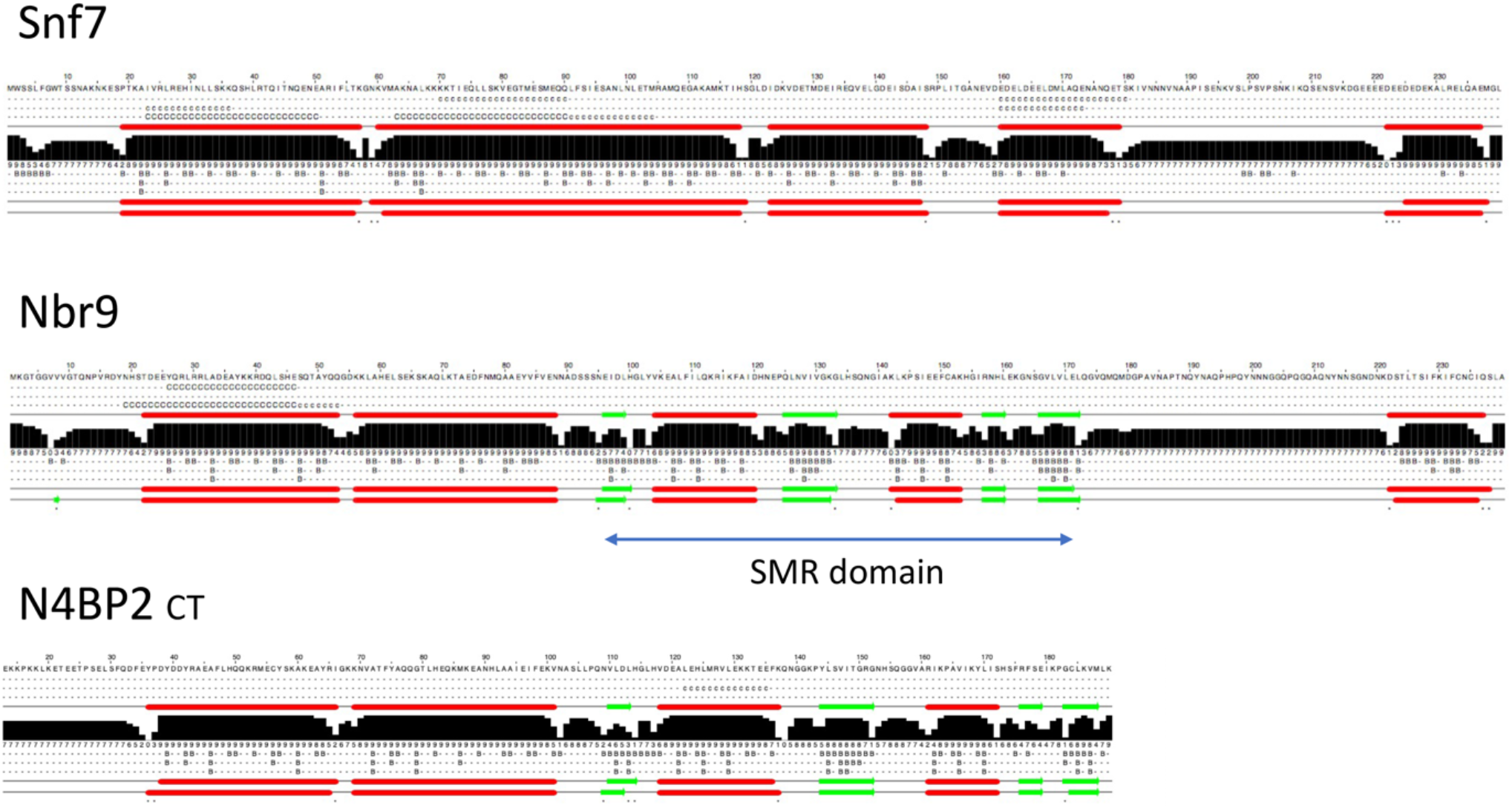
Comparison of the secondary structure of Nbr9 and Snf7. The secondary structures were determined by Jpred. Red bars: *α*-helical regions, green arrows: *β*-sheets. From top to bottom: Snf7, Nbr9 and the C-terminus of human N4BP2. The SMR domain in Nbr9 is marked.

**Table 2.** Comparison of Snf7 and Nbr9 *α*-helices.

| α-helix | Snf7<sup>1</sup> | Nbr9<sup>1</sup> |
| --- | --- | --- |
| 1 | 19-56 | 22-53 |
| 2 | 60-95 | 56-88 |
| 3 | 98-118 |  |
| 4 | 123-148 |  |
| 5 | 160-179 |  |
| 6 | 222-237 | 222-237 |

<sup>1</sup>amino acid position

This domain was originally found in bacterial MutS2 proteins thought to play a role in homologous recombination, but is now detected in almost all organisms. The SMR domain has endonuclease activity. There is a human homologue of Nbr9, Nedd4 binding protein 2 (N4BP2) (Murillas, Simms et al. 2002), whose C-terminal region is homologous to Nbr9 (Fig. 3). The function of this human protein is unknown. *NBR9* could be derived from a primordial ESCRT-III gene, where part of the coding region was replaced in a recombination event by SMR domain encoding sequences. This event must have occurred early in evolution, since these gene sequences are present in yeast and man.

To explore whether Snf7 and Nbr9 are structurally related, a structure prediction with AlphaFold was performed (Fig. 4). For Snf7 the predicted structure nicely matches the experimentally determined 3D-structure. Snf7 is displayed in its open conformation with a long extended *α*2-*α*3 helix (Tang, Henne et al. 2015). Further, AlphaFold predicts that the *α*6-helix aligns with *α*1. The position of the *α*6-helix could not be mapped in the available 3D-structures due to the high flexibility of the C-terminal region and because the ESCRT-III structures were generally made with C-terminally truncated proteins. For Nbr9 AlphaFold predicts the presence of an *α*1-*α*2 hairpin similar to Snf7 and interestingly, also an association between *α*6 and the *α*1-*α*2 hairpin, but here the interaction occurs with the *α*2-helix. Thus, there is indeed an overall similarity between the Snf7 and Nbr9 structures.

**Figure 4.**
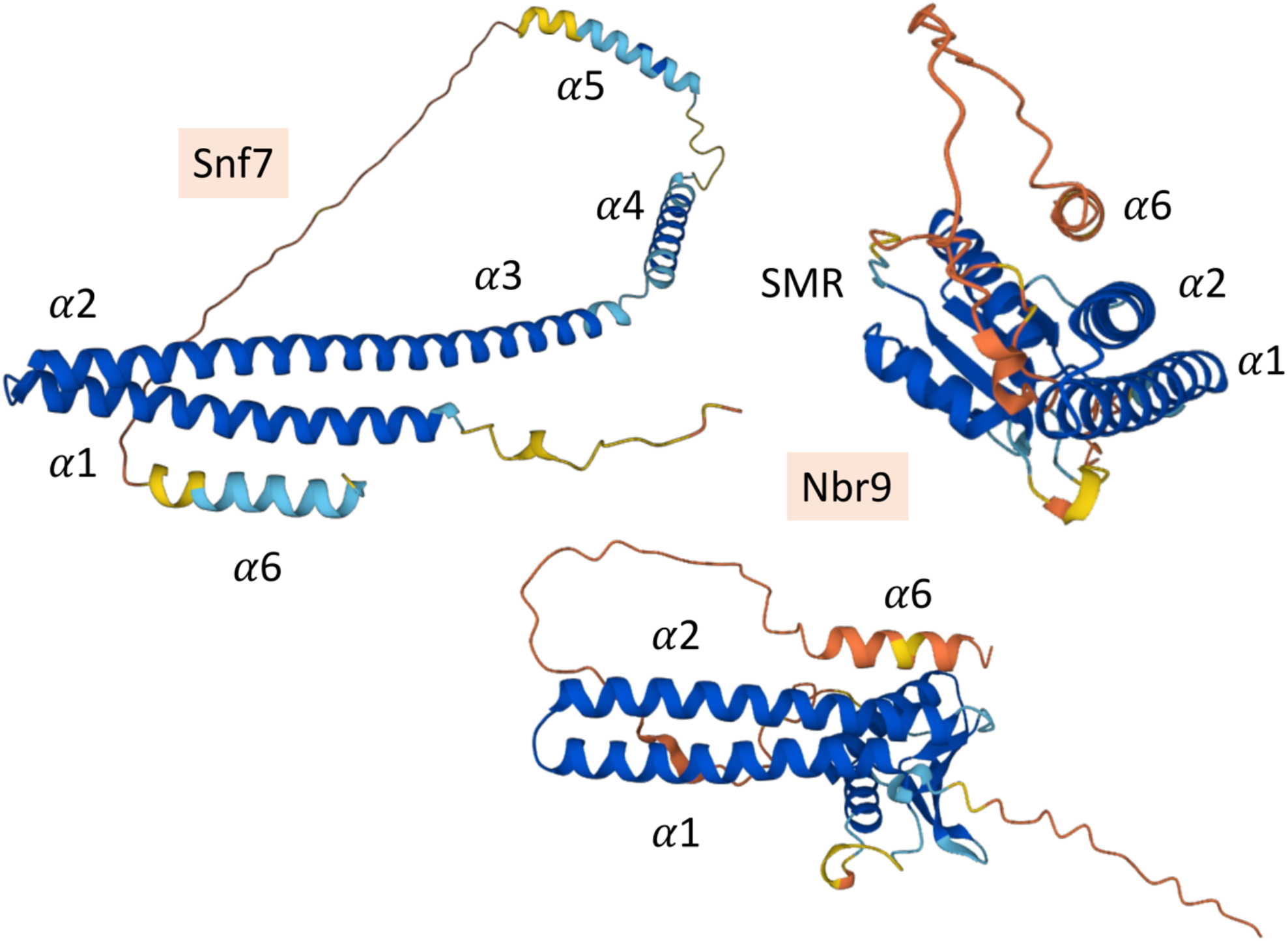
Structures of Snf7 and Nbr9 predicted by AlphaFold. Left: Snf7, right: Two views of Nbr9 rotated by 90°. The top view nicely shows the alignment of *α*6 with *α*2. Indicated are the Snf7 *α*-helices and the *α*-helices in Nbr9 corresponding to the helices in Snf7 (Tab. 2).

In the following we describe the outcome of various experiments that we performed with Nbr9. Most of the results were negative. Since it is not informative to show negative results, the figures of these negative experiments are not shown here. First, we were interested to see, whether Nbr9 shows ESCRT-III like behavior. Therefore, the *NBR9* deletion strain was examined for ESCRT-III knockout phenotypes. ESCRT-III deletion mutants show a number of growth phenotypes in the presence of different substances (Brune, Kunze-Schumacher et al. 2019), but none of compounds tested (caffeine, Congo Red, micafungin, LiCl, NaCl) had an effect on the growth of the *Δnbr9* strain. Also, unlike several of the ESCRT-III deletions, the *Δnbr9* strain was not temperature sensitive. Further, loss of Nbr9 did not stabilize the short-lived endocytic cargo protein Ste6 in cycloheximide chase experiments, thus Nbr9 is not involved in the degradation of endocytic cargo proteins via the multivesicular bodies (MVB) pathway. In this respect Nbr9 resembles Chm7 and Ist1, which are also not involved in MVB sorting (Bauer, Brune et al. 2015).

In addition, we were not able to detect an interaction with other ESCRT-III proteins in co-immunoprecipitation experiments. A similar negative result was also obtained for the established ESCRT-III protein Chm7. It appears that the Chm7 containing ESCRT-III complex only forms under specific conditions, which do not exist in exponentially growing haploid cells in rich medium at ambient temperature. Likewise, an Nbr9 containing complex could only form under special, so far unknown conditions. When *NBR9* was deleted in the octuple ESCRT-III deletion background, the yeast cells were still viable. Thus, so far, we do not have evidence that Nbr9 engages in the formation of an alternative ESCRT-III complex.

Nbr9 has been detected in two genome-wide screens. In one screen, it could be demonstrated that Nbr9 is palmitoylated in its C-terminal *α*-helix (Roth, Wan et al. 2006). To determine its intracellular localization, a Nbr9-ymNeongreen fusion, expressed from the chromosomal locus, was examined by fluorescence microscopy. With this fusion a clear staining of the plasma membrane could be observed (Fig. 5A). Inactivation of the palmitoylation site shifted the localization of the protein to the cytosol, demonstrating that the plasma membrane localization is mediated by palmitoylation. Staining was excluded from the vacuoles, but not from the nucleus.

**Figure 5.**
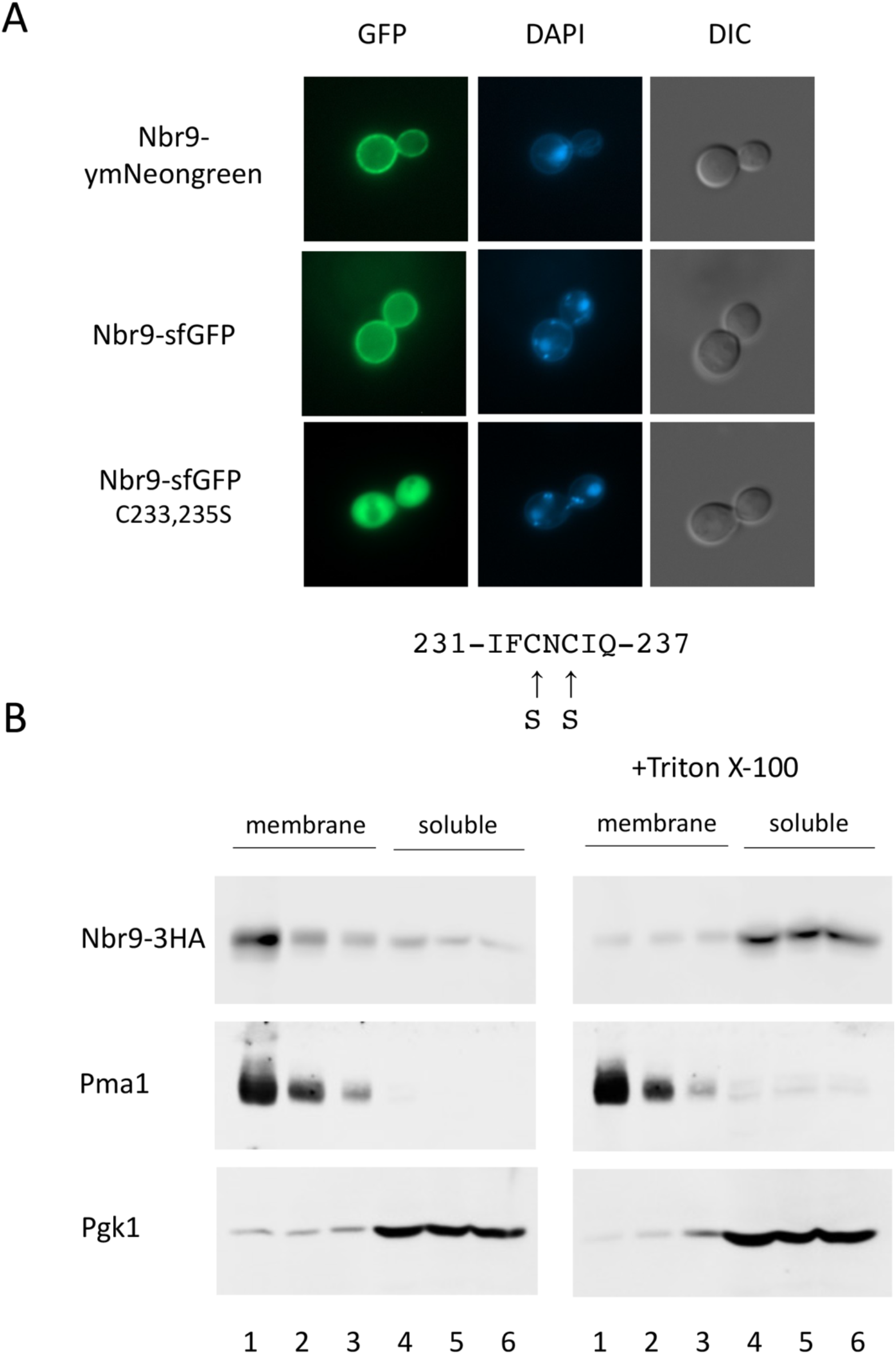
Localization and membrane association of Nbr9. (A) The localization of Nbr9 fusions was examined by fluorescence microscopy. From top to bottom: RKY3417 with an Nbr9-ymNeongreen fusion integrated at the chromosomal locus of *NBR9*, JD52 transformed with plasmid pRK1883 expressing an Nbr9-sfGFP fusion, JD52 transformed with plasmid pRK1891 expressing Nbr9-sfGFP with a mutated palmitoylation site as indicated at the bottom of the diagram. From left to right: GFP image, DAPI staining of the nuclei, DIC image. (B) Membrane association of Nbr9 is sensitive to Triton X-100 extraction. The membrane association of Nbr9-3HA was examined by a flotation experiment. The top three fractions of the gradients (lanes 1-3) contain the membranes, the lower three fractions (lanes 4-6) contain the soluble proteins. Cell extracts of strain RKY3021 were either treated (right panels) or not treated (left panels) with Triton X-100 before loading onto the gradients. Different proteins were detected by western blotting with specific antibodies. From top to bottom: Nbr9-3HA, Pma1 (raft associated membrane protein), Pgk1 (soluble protein).

Due to the presence of the SMR domain, Nbr9 is a putative nuclease. For the human homologue N4BP2, nuclease activity of the SMR domain has been reported (Watanabe, Wachi et al. 2003; Diercks, Ab et al. 2008; Jeong, Kim et al. 2012). We were not able to reproduce these results with yeast Nbr9, neither for the total protein nor for the isolated SMR domain. Although Nbr9-6His purified from *E. coli* displayed plasmid nicking and RNA degrading activity, the same activity was obtained with Mos10-6His used as a negative control. Thus, apparently, the nuclease activity was an unspecific contamination of our 6His purification. Nevertheless, it appears plausible that Nbr9 is a nuclease.

If Nbr9 is indeed a nuclease, it is difficult to envision what the function of the protein at the plasma membrane might be. Since palmitoylation is a reversible modification (Duncan and Gilman 1998), it seems more likely that the protein is sequestered at the plasma membrane and that it is released under specific circumstances to gain access to its target(s).

The bacterial MutS2 proteins have been implicated in DNA recombination. We therefore tested, if treatment with DNA damaging agents would cause a re-localization of Nbr9-sfGFP from the plasma membrane to the nucleus. But treatment with a number of DNA damaging agents (bleomycin, methyl-methane, sulfonate, hydroxyurea) for extended periods of time (up to 6 h) did not lead to a re-localization of Nbr9-sfGFP.

It has been suggested that Nbr9 could be involved in “Nonstop Decay” or “No Go Decay” of mRNA at stalling ribosomes (Glover, Burroughs et al. 2020). There are two proteins in yeast with an SMR domain. For one of these proteins, Cue2, a role in No Go Decay has been clearly established (D’Orazio, Wu et al. 2019). In analogy, it has been concluded that Nbr9 may be involved in mRNA decay as well (Glover, Burroughs et al. 2020). But a detailed analysis of the contribution of Nbr9 to mRNA decay is still lacking. It has been speculated that the ubiquitin binding CUE domains of Cue2 are important for binding to the ribosome (D’Orazio, Wu et al. 2019). Nbr9 does not have CUE domains, thus it would not be able to interact with ribosomes in the same way as Cue2.

*NBR9* was also detected in a visual screen for factors involved in cargo sorting to the plasma membrane (Proszynski, Klemm et al. 2005). In this screen, mutants from the yeast deletion library were screened for mis-sorting of the raft-associated Fus1-Mid2-GFP reporter protein. This protein is normally transported to the plasma membrane. In some mutants, among them *Δnbr9*, it was mis-localized to internal structures or to the vacuole. However, we were not able to reproduce this phenotype. In our *NBR9* deletion strain, the localization of Fus1-Mid2-GFP was not altered compared to wildtype.

It has been suggested that palmitoylation promotes association with sphingolipid- and ergosterol-rich lipid rafts (Levental, Lingwood et al. 2010). To test, whether the palmitoylated Nbr9 protein as well is associated with rafts, a flotation experiment was performed (Fig. 5B). Lipid rafts cannot be dissolved by treatment with the detergent Triton X-100 (Bagnat, Keranen et al. 2000), because of this property, they are also called “detergent resistant membranes” (DRMs). This can be clearly seen for the raft-associated Pma1 protein, which was resistant to detergent extraction and stayed in the membrane fraction after Triton X-100 treatment. But, Nbr9-3HA in contrast was shifted from the floating membrane fraction to the soluble fraction by Triton X-100. Thus, Nbr9 does not appear to be associated with lipid rafts.

To summarize, several yeast proteins were detected in this study with similarities to the ESCRT-III protein Snf7. Whether these proteins only share an ancient protein motif with ESCRT-III proteins or whether they are indeed distant members of the ESCRT-III protein family remains to be established. The Nbr9 protein strikingly resembles Snf7 in its secondary structure. Therefore, chances are high that Nbr9 is indeed a true member of the ESCRT-III family.

## Compliance with Ethical Standards

This study did not involve human participants or animals.

## Supplement

**Figure S1.**
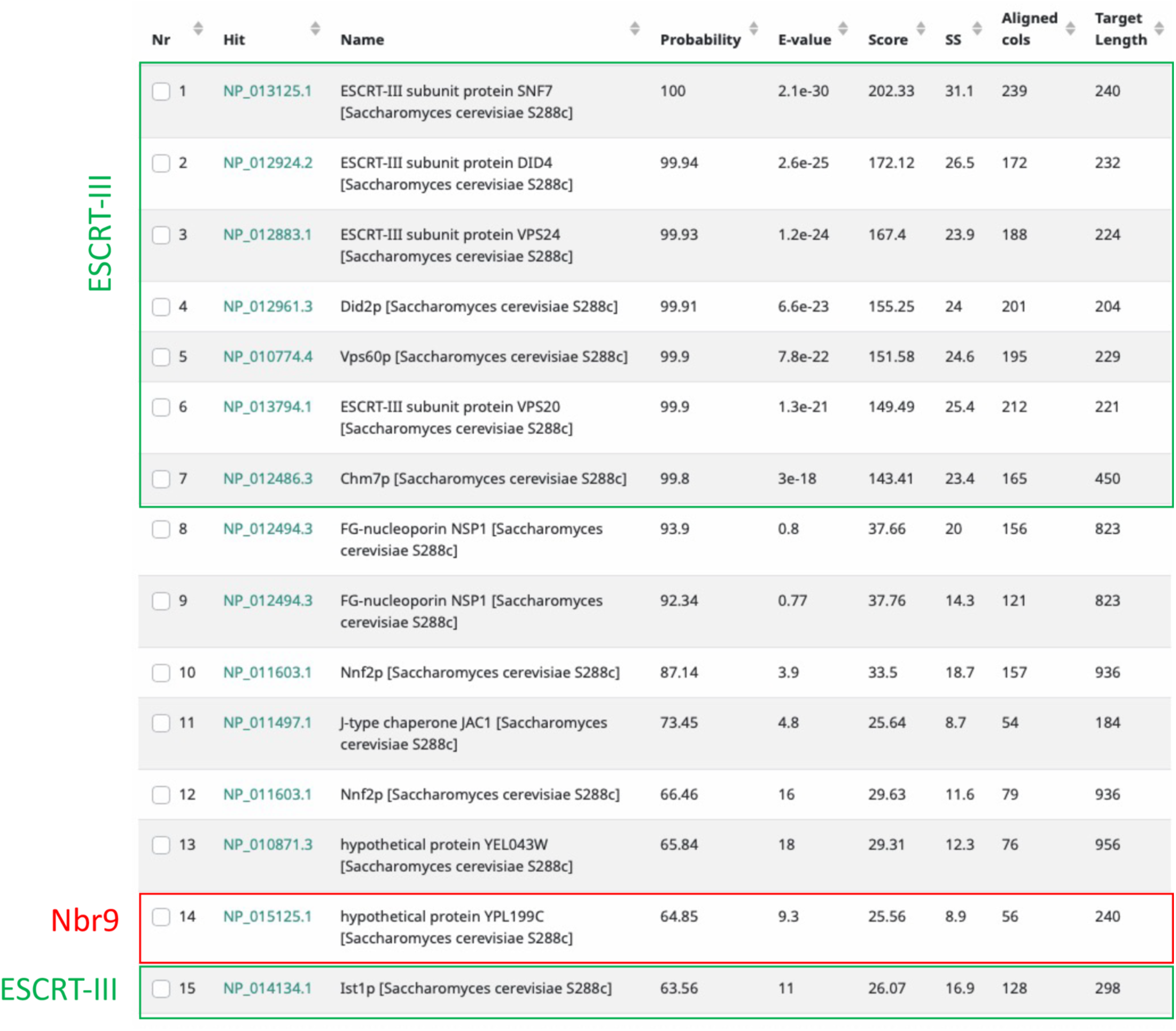
HHPred output with Snf7 as a query. The first 15 hits are shown. Green boxes: ESCRT-III proteins, red box: Nbr9.

**Figure S2.**
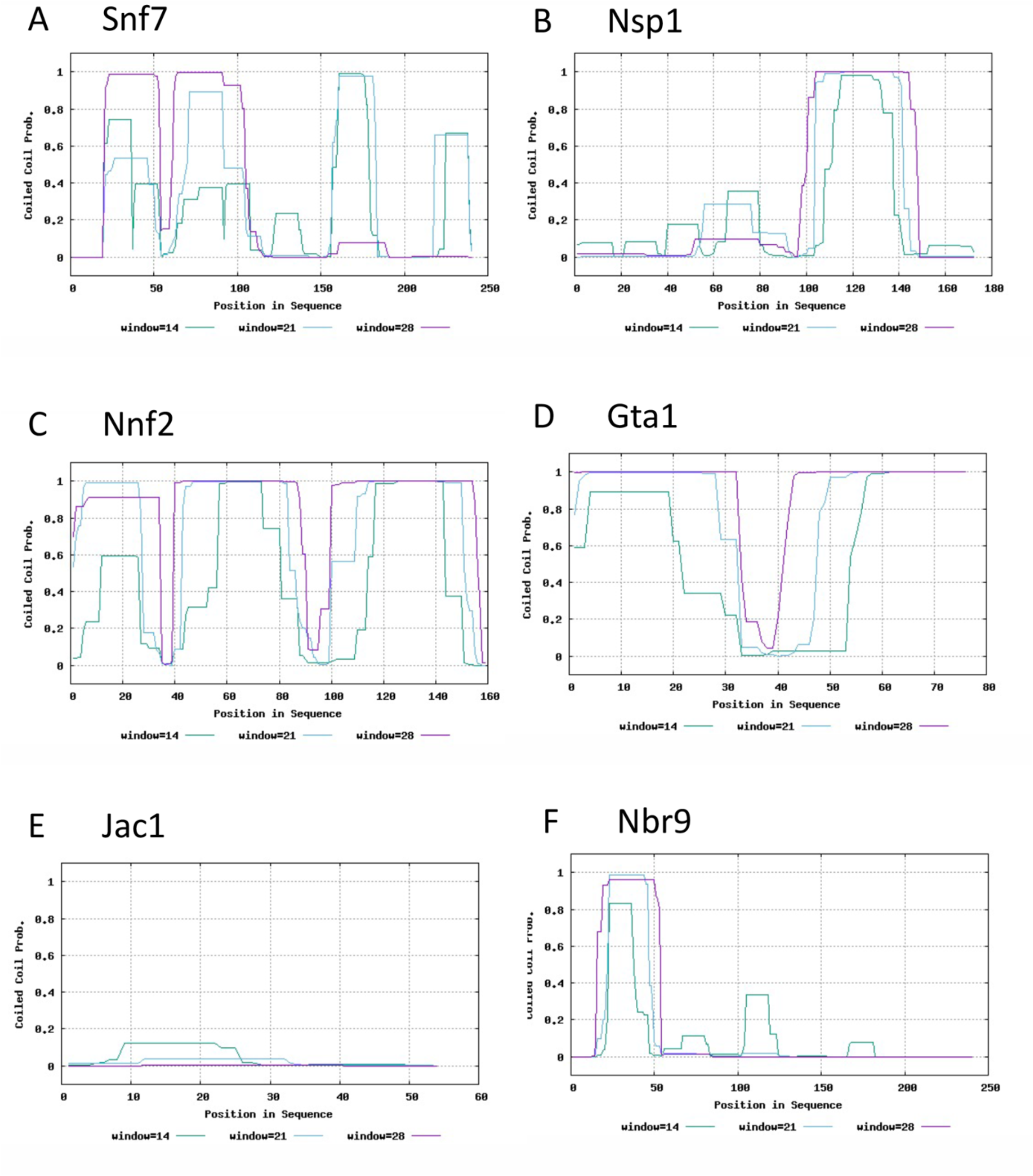
Prediction of coiled-coils with PCOIL for the protein regions similar to Snf7. Y-axis: coiled-coil probability with different window sizes (14, 21, 28 amino acids).

**Figure S3.**
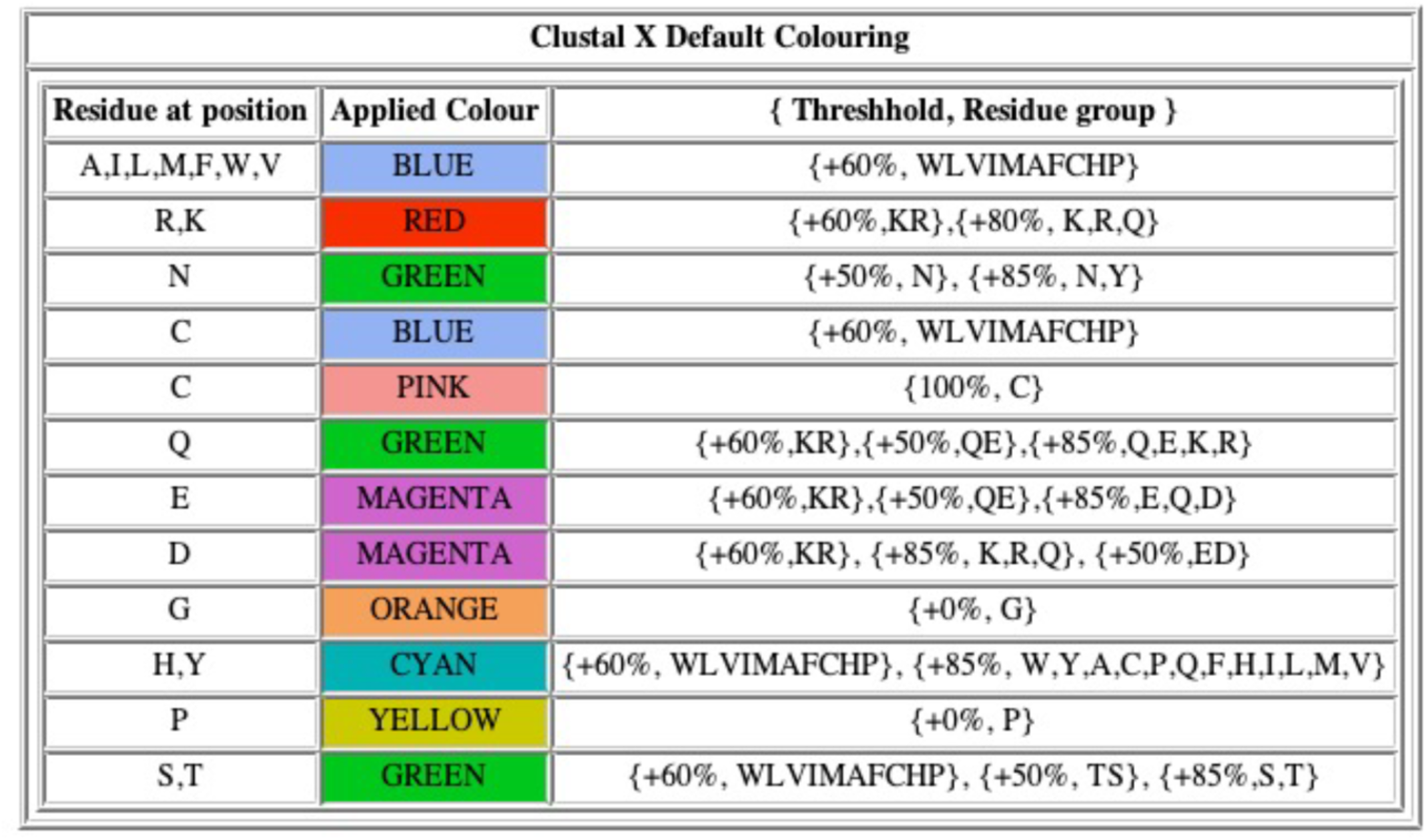
Clustal X default coloring scheme used for Fig. 2.

